# Systematic Mapping of cell Wall Mechanics in the Regulation of Cell Morphogenesis

**DOI:** 10.1101/502229

**Authors:** Valeria Davì, Haotian Guo, Hirokazu Tanimoto, Katia Barett, Etienne Couturier, Arezki Boudaoud, Nicolas Minc

## Abstract

Walled cells of plants, fungi and bacteria, come with a large range of shapes and sizes, which are ultimately dictated by the mechanics of their cell wall. This stiff and thin polymeric layer encases the plasma membrane and protects the cells mechanically by opposing large turgor pressure derived stresses. To date, however, we still lack a quantitative understanding for how local and/or global mechanical properties of the wall support cell morphogenesis. Here, we combine super-resolution imaging, and laser-mediated wall relaxation, to quantitate subcellular values of wall thickness (h) and bulk elastic moduli (Y) in large populations of live mutant cells and conditions affecting cell diameter in the rod-shaped model fission yeast. We find that lateral wall stiffness, defined by the surface modulus, σ=hY, robustly scales with cell diameters. This scaling is valid in tens of mutants covering various functions, within the population of individual isogenic strains, along single misshaped cells, and even across the fission yeasts clade. Dynamic modulations of cell diameter by chemical and/or mechanical means suggest that the cell wall can rapidly adapt its surface mechanics, rendering stretched wall portions stiffer than unstretched ones. Size-dependent wall stiffening constrains diameter definition and limits size variations, and may also provide and efficient mean to keep elastic strains in the wall below failure strains potentially promoting cell survival. This quantitative set of data impacts our current understanding of the mechanics of cell walls, and its contribution to morphogenesis.

## Introduction

The cell wall (CW) is a ubiquitous structural layer defining the surface properties of most microbial and plant cells (1-3). It is composed of long cross-linked sugar strands and proteins, which form a polymeric matrix with a thickness of few tens up to few hundreds of nanometers. The mechanical properties of the CW are largely determined by its thickness and geometry, and by the composition and arrangement of its polymeric network, which impinge on material anisotropy and bulk elasticity (4–6). Those properties are patterned from the biochemistry of wall synthesis and remodeling, which is spatially and temporally regulated during cell growth (5). At short time-scales, the cell wall exhibits elastic-like properties and is put under tension by a large internal turgor pressure typical of walled cells (7–9). As such, the balance between turgor values and wall stiffness has long been predicted to influence the instantaneous shapes of walled cells (10–12). Given the large range of shapes and sizes spanned by walled cells, addressing mechanical strategies and dose-dependent modulations of cell wall properties remain an outstanding open problem in morphogenesis.

Measuring the mechanical properties of the cell wall is a long-term endeavor in plant and microbial sciences. Methods to measure wall stiffness include Atomic Force Microscopy (AFM) (13), active cell bending or compression (7, 8, 14), and strain-stress assays through the modulation of turgor pressure (10, 11, 15–18). Those allow to extract a surface elasticity but are in general low-throughput, and importantly do not take into account putative global or local variations in wall thickness, thereby preventing to discern contributions from bulk material properties vs thickness to cell wall elasticity. This has limited our appreciation of wall mechanical and biochemical regulation, as bulk elasticity likely reflects polymer composition and cross-linking, while thickness is expected to depend on a balance between synthesis and deformation rates (5, 16). In addition, those individual parameters have a differential impact on the stress, elastic strain or bending modulus of the wall (12). Therefore, addressing how individual mechanical parameters locally and globally vary with cell shape and size (19) is essential for a complete conceptual understanding of the contribution of wall mechanics to morphogenesis.

In here, we combined sub-resolution imaging of CW thickness and laser-mediated wall relaxation assays to systematically map bulk elasticity and thickness, with subcellular resolution and medium-throughput. We use this approach to address how mechanical properties of the lateral CW of rod-shaped fission yeast cells vary with cell diameter. Using libraries of mutants defective in diameter regulation, we find that the wall is systematically stiffer in wider cells, and in wider subcellular regions of mis-shaped cells. We evidence a wall stiffening mechanism, associated with dynamic diameter increase, in which wall sections become gradually stiffer as they are deformed laterally. Size-dependent wall stiffening maintains elastic strains in the wall to low and near-constant values and constrains diameter definition, with crucial implications for cell viability and size regulation.

## Results

### A medium-throughput method to map subcellular mechanical properties of the cell wall

The fission yeast *Schizosaccaromyces pombe* is one of the most-established model to understand the emergence of cell morphogenesis (20–23). Those are rod-shaped cells which grow by tip extension with a near-constant diameter of ~4 microns. As for other walled cells, their shape is defined by their CW encasing the plasma membrane (5). Building on previous methodologies to map CW thickness and compute its surface stiffness (10, 11, 15, 16), we sought to develop a systematic approach to map those key mechanical parameters in large populations of cycling cells and mutants with medium-throughput (Fig. 1A). We constructed strains expressing GFP-Psy1, to label the plasma membrane at the inner side of the wall, and added lectins from *Griffonia simplicifolia* tagged with a far-red fluorophore (AlexaFluor-647), to mark the outer face of the wall. As previously reported, this, combined with a dedicated image analysis pipeline, allows to map CW thickness around a given cell, as well as to delineate CW shape with nanometric resolution (16) (Fig. 1B). After imaging a cell to compute wall thickness, we immediately deflated it by piercing the wall with a UV laser focused at a small diffraction-limited point on the surface (11, 15, 24). Cells deflated with a marked and homogenous reduction in the lateral radius, as well as changes in curvature at cell tips. Cell shape analysis before and less than 1 min after ablation allowed to compute the local value of the ratio of the surface modulus of the CW to turgor pressure, hY/P (see material and methods) (11) (Fig. 1C). The combination of both methods, thus served to independently compute wall bulk elastic moduli (Y) and thickness (h), and test their contribution to morphogenesis.

**Figure 1.**
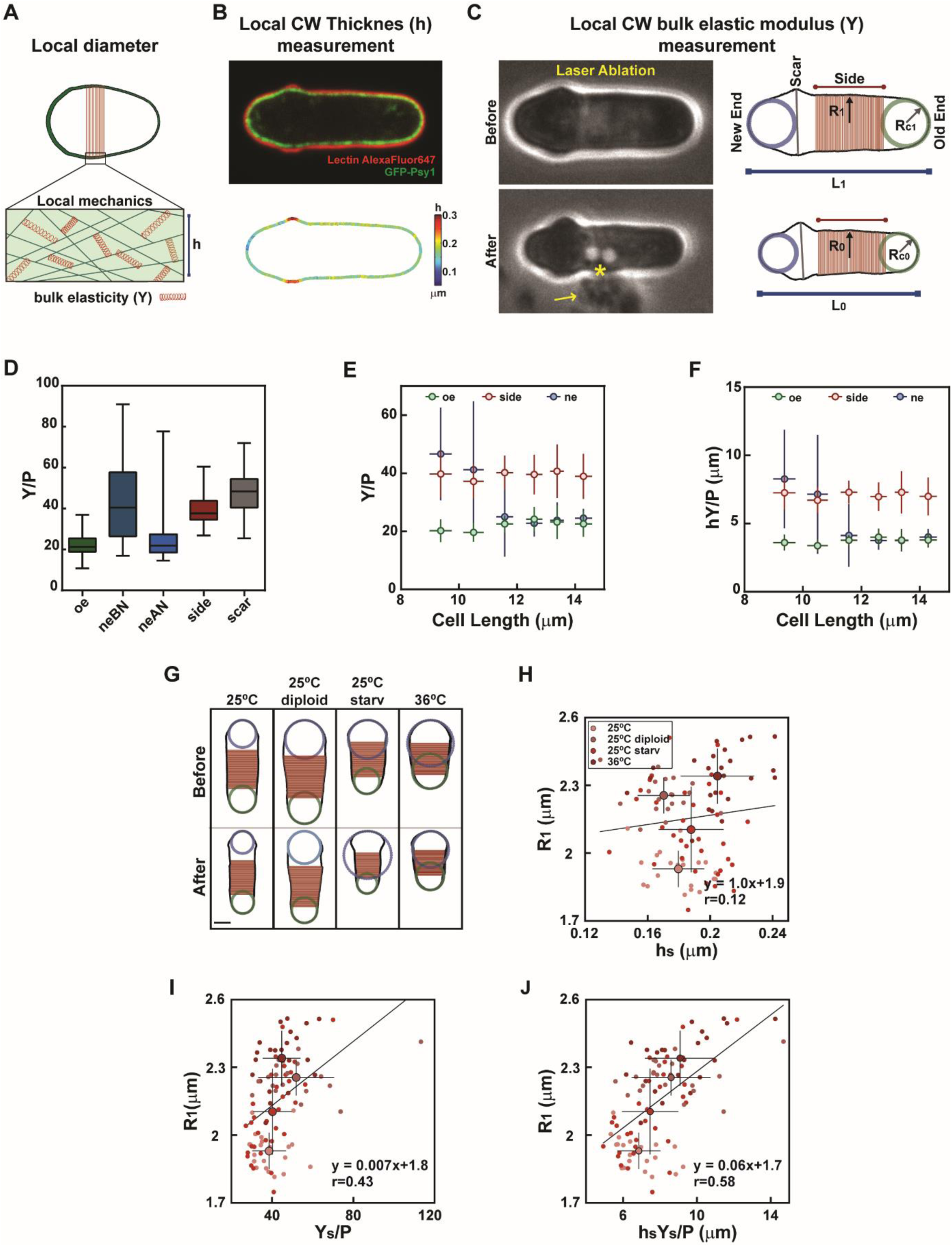
A method to compute local cell wall mechanical properties in *S. pombe* cells. **(A)** Schematic representation of a cell with varying diameters, and zoom on local cell wall organization, defining thickness (h) and bulk elastic modulus (Y). **(B)** Mid-slice confocal image of a cell expressing GFP-Psy1 and labelled with lectin-Alexafluor647, and corresponding CW thickness map obtained as in (16). **(C)** (Left) Method to estimate local wall bulk modulus (Y) divided by turgor pressure (P). Bright-field images of the same cell depicted in B before (top) and after (bottom) laser ablation. The star indicates the location of laser ablation, the arrow point at a cloud of cytosolic material ejected from the cell. (Rigth) Cell wall boundaries of the same cell, obtained by binarization of the lectin-Alexafluor647 signal, before and after ablation, used to compute local radii of curvature at cell tips, and the radius along the cell side, defined as a mean on the red region. **(D)** Y/P values computed at old end (oe, n=99), new end before (neBN, n=20) and after (neAN, n=64)) NETO, sides (n=99) and scars (n=27), in a WT population. Evolution of Y/P **(E)** and hY/P **(F)** as a function of cell length. Individual cells have been imaged and binned by length, (between 8 and 20 cells for each point). **(G)** Cell wall boundaries and local shape features, before and after ablation, of typical wt haploid and diploid cells grown at 25°C in exponential phase (respectively 25°C and 25°C diploid), and haploid cells after 16 hs of starvation (25°C starv) and at 36°C in exponential phase (36°C). **(H-J)** Cell radius (R_1_), as a function of side wall thickness (h_s_), side bulk modulus divided by pressure (Y_s_/P) or surface modulus divided by P (Y_s_h_s_/P), in the same conditions as in F (for 25°C n=21, 25°C diploid n= 19, 25°C starv n=34, 36°C n=24). Small dots correspond to single cells measurements, and larger dots represent the average values for each condition. The line is a linear fit on single cell measurements. r values correspond to Pearson correlation coefficients. Whisker plots represent median and full data set range. Error bars are standard deviations. Scale bars, 2μm.

In wild-type cells, this approach yielded near similar values of wall anisotropic extension as in previous reports (Fig. S1A-S1B) (11), and similar mean values of bulk elastic moduli of ~ 50 MPa using previous estimates of turgor pressure of 1.5 MPa (7, 10, 11, 16). We first explored subcellular patterns of wall mechanics throughout the cell cycle. The growing old end and new end after New End Take Off (NETO) were the softer parts of the cell, likely accounting for growth and wall remodeling there. The birth scars, cell sides and non-growing new ends had a bulk elasticity typically ~2X higher than growing ends (Fig. 1D and Fig. S1C). Sorting cells by length revealed that the old end and cell sides kept near-constant bulk and surface elastic moduli throughout the cell cycle. In contrast, the new end underwent a marked two-fold reduction in bulk and surface moduli at a length of around 10–12 µm, likely corresponding to growth resumption there after NETO (Fig.1E and Fig. S1C).

Using WT populations, we next asked if local cell shape features were correlated with specific CW mechanical parameters around the cell. Because normal haploid WT cells have a narrow range of diameters, we used starved cells, cells grown at 36°C, and diploids, conditions known to affect cell diameters. This analysis showed that the local radius of curvature at the growing old end is only mildly correlated with CW thickness, elastic and surface moduli divided by pressure. Similarly, radii at non-growing new ends exhibited weak positive correlation with their local counterpart values of bulk and surface moduli, and almost no correlation with thickness (Fig. S1D-S1J). However, the radius along the cylinder, exhibited the highest positive correlation with side surface moduli, indicating that this mechanical parameter could encode for diameter regulation (Fig. 1G-1J and Fig. S1J).

### The surface modulus of the cell wall scales with cell diameter

Building on those observations, we asked if mechanical properties of the CW would vary in populations of mutants with defective diameter regulation. For this, we examined 26 mutants knocked out for a single open reading frame, which have been reported to bear defects in cell diameter from previous visual screens of the fission yeast KO library (22, 23). After careful validation and diameter quantification (see material and methods), we refocused our analysis, on a specific set of 18 mutants, which solely exhibited diameter mis-regulation, discarding more complex misshaped mutants with other bent or branched features, for instance (22, 23). Importantly, those mutants covered several physiological processes, including cell polarity, chromosome segregation, signaling, metabolism and wall synthesis (Table 1, Fig. 2A-E and Fig. S2A). Defects in diameter regulation in those strains could be segregated into 3 sub-categories. One category had a mean diameter significantly different (higher or lower) than WT (Fig. 2A and 2C). A second category had a similar mean value as WT diameters, but a much larger variability (computed as a standard deviation), reflecting defects in diameter maintenance through successive divisions (Fig. 2D). A last category was composed of skittle-shaped mutants with defects in diameter along a single cell (Fig. 2B). Finally, we added to this screen, 8 mutants involved in CW regulation, which do not have major defects in diameter, but are expected to mis-regulate wall mechanics. To compute wall mechanical parameters, mutant strains were transformed to stably express GFP-psy1, and grown and assayed as controls following a standardized procedure. For each strain, we typically analyzed ~ 20–30 cells, discarding dividing cells. In addition, although we kept mutants that may have skittle-like shapes, we first analyzed a sub-population in those strains with near intact rod-shapes.

**Figure 2.**
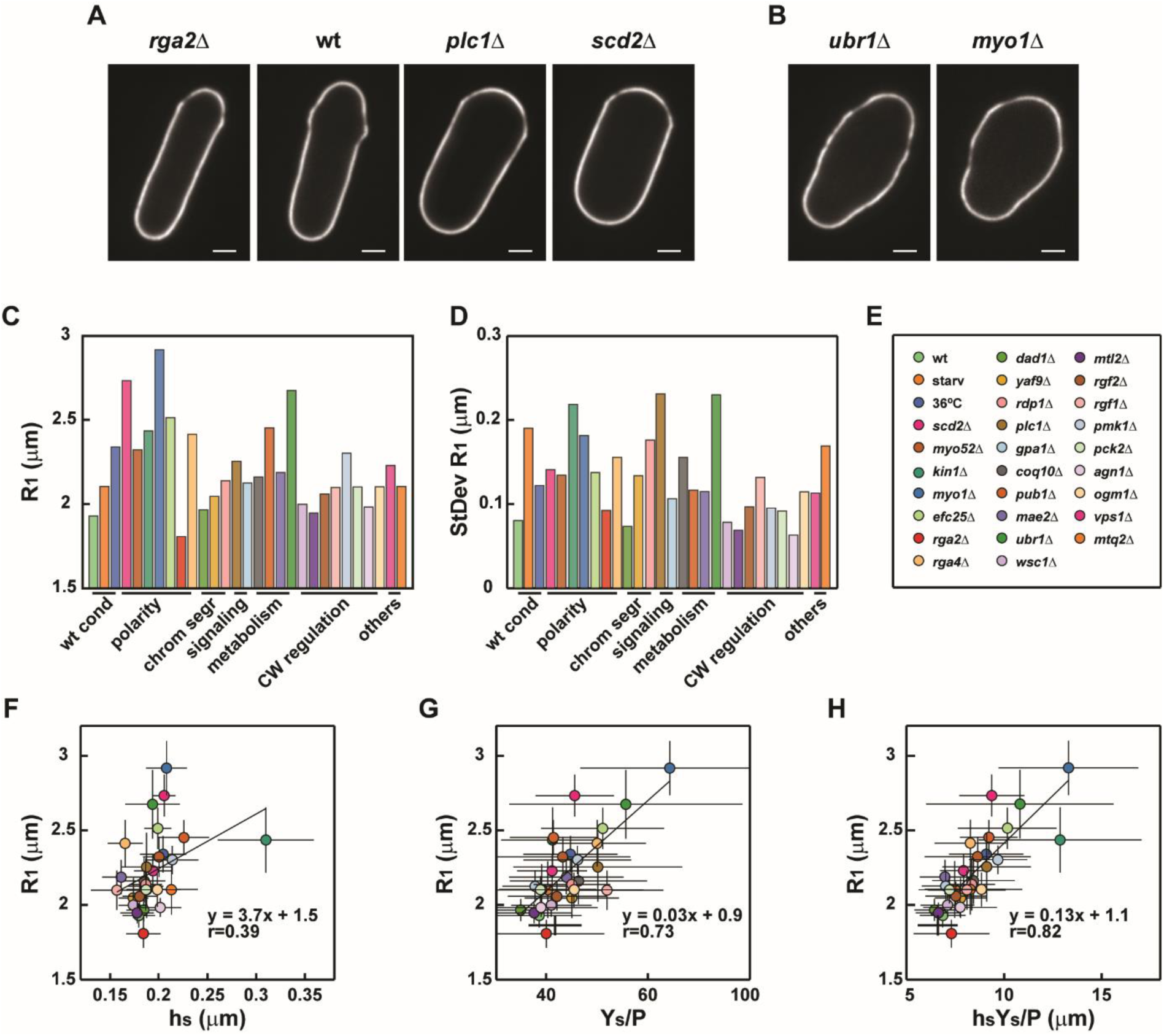
Mapping cell wall mechanics in populations of mutants defective in cell diameter. **(A-B)** Mid-slice confocal image of typical cells labelled with lectin-Alexafluor647 of wt and mutant strains, with defects in mean diameter (A) and diameter along their length (G) **(C-E)** Mean cell radius on cell sides of different mutants and conditions **(C)** and standard deviations **(D)** as annotated in the legend **(E)**. Mutants are sorted by a simplification of the Gene Ontology functions described in Table S1. **(F-H)** Cell radius (R_1_), plotted as a function of side wall thickness (h_s_), side bulk modulus divided by pressure (Y_s_/P) or side surface modulus divided by pressure (Y_s_h_s_/P), in the conditions and mutants listed in the legend **E**. Each strain is represented by its average value. The line is a linear fit on mean values for each condition. r values correspond to Pearson correlation coefficients. Scale bars, 2μm.

This analysis over tens of mutants, revealed a relatively narrow distribution of side CW thickness around 200nm, with one particular mutant, *kin1Δ*, departing from the main trend, and exhibiting a higher thickness of more than 300nm (25) (Fig. 2F). Correlation between diameter and thickness was positive, but relatively low (Pearson’s coefficient, r=0.39) (Fig 2F). Elastic moduli divided by pressure, Y/P, also increased with cell diameter, and displayed a much higher correlation (r=0.73) (Fig. 2G). Importantly, those variations in the values of Y/P mostly reflected changes in the bulk modulus of the wall and not turgor pressure. This was evidenced by comparing the relaxed length obtained from wall piercing through laser ablation, and the one obtained with increasing amounts of sorbitol treatment (11). This analysis performed in mutants with the largest diameters, yielded a relative pressure compared to that of WT cells, and revealed variations below few % (Fig. S2B-S2C). This suggests that mutants with varying diameters may have similar turgor values, and that shape variations may be mostly associated with changes in CW mechanics.

The surface modulus, exhibited the highest correlation coefficient with the cell radius (r=0.82) (Fig. 2H). Accordingly, mutants which departed from the main trend in thickness or bulk elastic modulus appeared to compensate the other parameter, so that the surface modulus remained in the mean trend. For instance, *kin1Δ*, exhibited a much larger thickness than mutants with similar diameters, but also featured a reduced bulk modulus yielding a surface modulus close to the main trend (Fig. 2F-2H). We also found a good correlation between thickness, bulk and surface moduli at the old end to that at cell sides, suggesting that mechanical properties of the lateral CW could be defined during the initial synthesis of the CW at cell tips and modified in the same manner during growth. In contrast, the mechanical properties of the CW at the new end were much less correlated with that of the sides, plausibly as a consequence of prior septum synthesis there (11) (Fig. S2H-S2J). Thus, this large-scale analysis, suggest a dose-dependent link between cell diameter and CW surface modulus, which best represents the stiffness of this structure.

### Size-dependent wall stiffening in isogenic populations, at subcellular levels, and across the fission yeast clade

We then tested if wall mechanics and diameters were correlated between single cells in an isogenic population. For this, we re-examined a mutant strain, *plc1Δ*, deleted for a gene which encodes for a phospholipase C enzyme, and exhibits rod-shaped cells with the highest standard deviation in diameter (Fig. 2A-2E) (26). When analyzed at the level of a population of single mutant cells, we found a net positive correlation between CW surface modulus and cell diameter (r=0.62) (Fig. 3A). Importantly, this correlation was mostly associated with an increase in bulk modulus of the wall of fatter cells (Fig. S3A-S3B). Thus, modulation of cell wall stiffness is associated with cell to cell variations in diameter.

**Figure 3.**
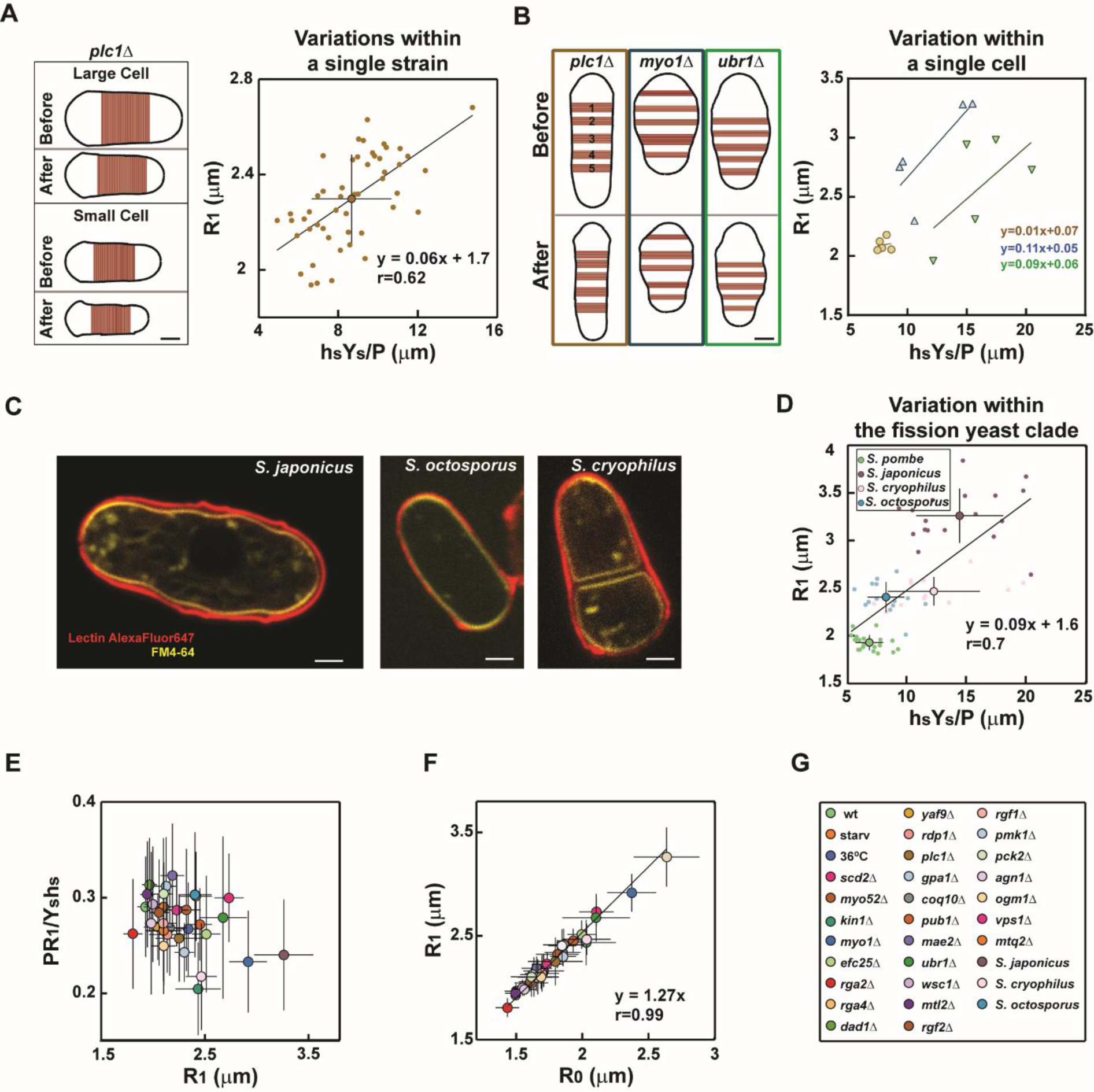
Estimation of cell wall mechanical parameters in a single isogenic strain, along single mis-shaped cells and in other fission yeast species. **(A)** (Left) Cell Wall boundaries, before and after ablation, of 2 extreme *plc1Δ* cell examples with large and small radii. (Right) Cell radius (R_1_) plotted as a function of the surface modulus divided by P (Y_s_h_s_/P) within a population of *plc1Δ* cells (n=50). **(B)** (Left) Cell wall boundaries and definition of 5 cell wall portions of cell sides, before and after ablation, for one rod shape cell (*plc1Δ*) and two skittles cells (*myo1Δ* and *ubr1Δ*). (Right) Local radius (R_1_), plotted as a function of the corresponding surface modulus divided by P (Y_s_h_s_/P) in 5 different parts along the side of the same cell, for the same cells represented on the left (colors correspond). **(C)** Representative mid-slice confocal images of a cell from each fission yeast species labelled with FM4–64 (membrane) and lectin-Alexafluor647 (wall surface). **(D)** Cell radius (R_1_) plotted as a function of side wall surface modulus divided by P (Y_s_h_s_/P) for *S. pombe* (n=21), S. *japonicus* (n=17), *S. octosporus* (n=14) and *S. cryophylus* (n=12) **(E)** Elastic strain in the cell wall (PR_1_/ Y_s_h_s_) plotted as a function of cell radius **(F)** Turgid radius, R_1_ plotted as a function of the relaxed radius, R_0_ in the conditions, mutants and species listed in **(G)** legend. In A-D lines in plots represent linear fits of single cell measurements. In E-F the line is a linear fit of mean values for each condition. r corresponds to Pearson correlation coefficients. Error bars represent standard deviations, scale bars 2μm.

To understand if the positive correlation between cell diameter and wall stiffness was also valid at a local level, we analyzed skittle-shaped cells. We selected 2 representative mutants with a high penetrance of skittle-like defects, but pertaining to distinct genotypic classes (*myo1Δ* and *ubr1Δ*), and used the *plc1Δ* rod-like cells, as controls. We computed local diameters along the cell long axis in one representative cell from each mutant, and plotted them as a function of local surface moduli. While points in the rod-shaped *plc1Δ* cell clustered around a single value for the radius and surface modulus, those parameters varied and were strongly correlated along the length of single *myo1Δ* and *ubr1Δ* cells, with larger portions of CW being stiffer (Fig. 3B and Fig. S3C-S3D). These local estimations were also used to simulate inflation of relaxed walls, yielding to near-similar original cell shapes, confirming the accuracy of our method used in sections of non-perfect rods such as skittle cells (Fig. S3E). Those local effects, further rule out putative contributions from turgor variations, and strongly suggest that the surface modulus of the CW, provides a robust predictor for local as well as global diameter values in fission yeast.

We next tested if correlations between wall stiffness and diameter, could be a conserved feature in evolution. We measured CW mechanical properties in closely related species pertaining to the fission yeast clade, *Schizosaccaromyces japonicus*, *Schizosaccaromyces octoporus* and *Schizosaccaromyces cryophilus*. Those species exhibit cells with markedly different diameters, as compared to *S. pombe* cells. Remarkably, the scaling between the surface moduli and diameters was also highly pronounced in this clade (Fig. 3C). However, in contrast to *S. pombe* mutants, variations in stiffness were mostly associated to thickness changes of the lateral wall, with some large *S. japonicus* cells featuring walls typically twice as thick as *S. pombe* cells (Fig. S3I-S3J). This suggests that bulk material properties and plausibly compositions of walls in those different species, may not drastically vary. Rather, adaptation of wall thickness appears as the dominant strategy to scale wall stiffness to cell size in the fission yeast clade.

### Size-dependent lateral wall stiffness limits elastic strains

Using aforementioned data, we then computed which conceptual mechanical parameter may constrain cell diameter definition. We computed the stress born by the lateral CW, defined as PR/h, the elastic strains defined as PR/Yh, and the bending energy of the CW normalized by that of pressure, computed as PR^3^/Yh^3^. Without size-dependent adaptation in thickness, bulk elasticity or pressure, those parameters are expected to steadily increase with cell diameters. The stress in the CW estimated assuming constant values of pressure of 1.5 MPa systematically increased with diameter. The CW of fatter cells thus bears higher stresses than in smaller cells, suggesting that stress limitation may not be a strategy to control the shapes of those walled cells (19) (Fig S3H). The bending energy was more variable, but also increased with diameter, suggesting that CW bending may not constrain diameter definition (12) (Fig. S3I). Remarkably, however, we found that the elastic strain in the wall was mostly constant with values around ~25–30%, and even slightly decreasing at larger diameter values (Fig. 3D). Accordingly, the relaxed cell radius, R_0_, was strongly correlated with the pressurized radius R_1_ (r=0.99), so that R_1_~1.3R_0,_ in agreement with a near-constant elastic strain of ~30% (Fig. 3E). Thus, this analysis demonstrates that diameter-dependent cell wall stiffening limits elastic strains in the wall.

### Dynamic modulations of cell diameters reveal strain-stiffening properties of the lateral cell wall

Walled cells may undergo rapid shape or size changes. To understand how mechanical properties of CWs may be modified during those events, we used chemical and physical approaches to dynamically modulate cell diameter, and monitor wall mechanics. We first treated cells with a low dose (10 µM) of the actin-depolymerizing drug Latruncunlin A. This intermediate dose may weaken polar domains of the active-GTP-bound Cdc42, and yields fatter cells (27). By performing a time-course we noted that cells enlarged their diameters in times as short as 60–90 min, without building a whole new CW through growth and division (Fig. S4A-S4B). Rather, the glucan synthase Bgs4, exhibited a gradual dispersion from cell tips to the whole cell contour over ~1h, probably yielding ectopic CW synthesis and diffuse growth, thereby enlarging diameter (Fig. S4C-S4D). By computing thickness and bulk elastic moduli in cells treated with 10 µM LatA for 90min, we found that diameter increase was associated with a reduction of lateral wall thickness, concomitant with an increase in bulk elastic modulus (Fig. 4A-4B and Fig. S4E-S4F). Importantly, those changes resulted in a net increase of the surface modulus on cell sides, yielding a near-conservation of CW elastic strains during these dynamic changes in diameter (23% in LatA, as compared to 26% in DMSO controls).

**Figure 4.**
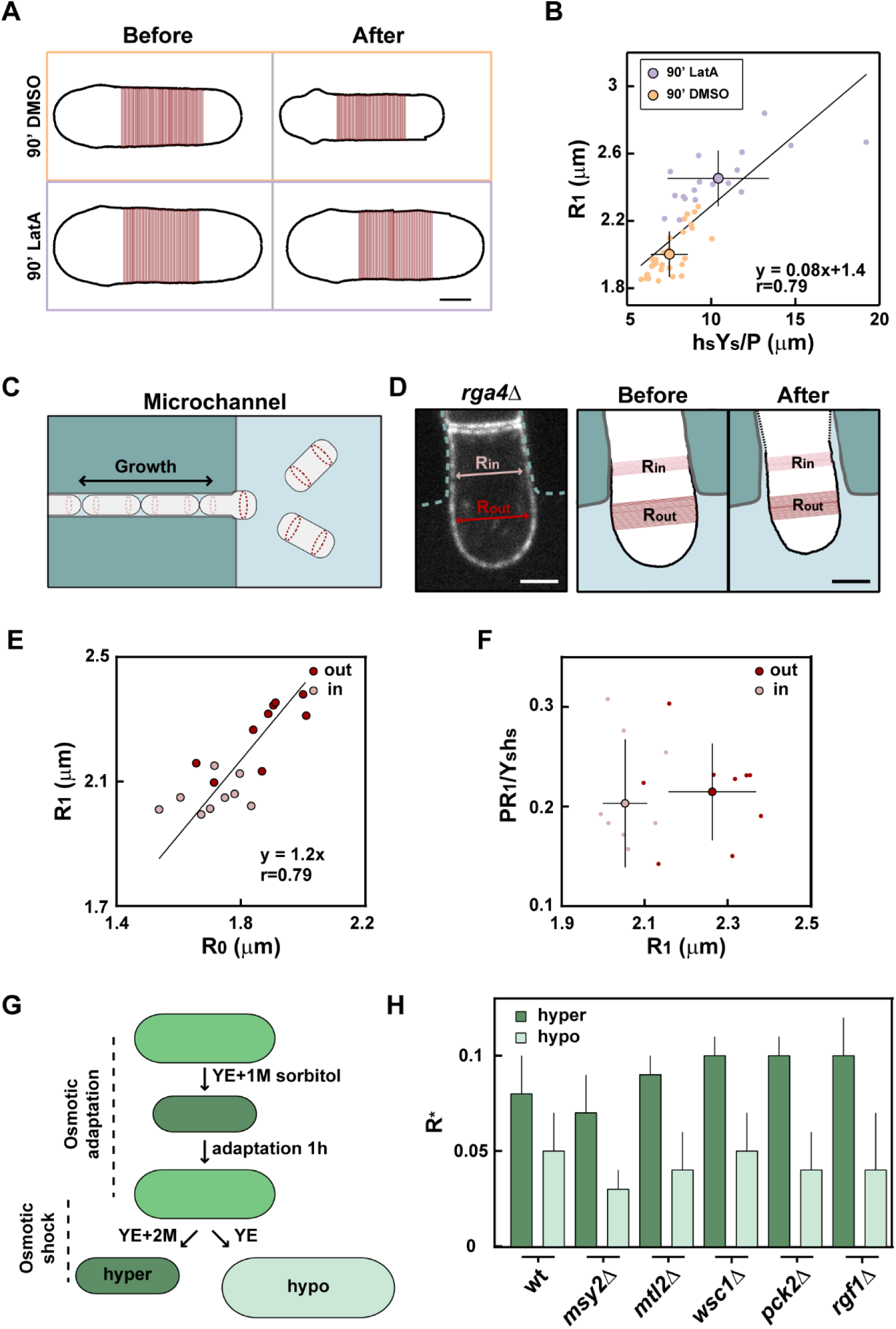
Dynamic modulations of CW mechanical parameters during local and global shape changes. **(A)** Cell wall boundaries and cell side definition, before and after ablation, of typical cells treated for 90’ with DMSO or 10μM Latrunculin A **(B)** Cell radius (R_1_), plotted as a function of side wall surface modulus divided by P (Y_s_h_s_/P), in wt cells treated for 90’ with DMSO (n=30) or 10μM LatrunculinA (n=18). Small dots are single cells measurements, and larger dots are averages. The line is a linear fit on single cell measurements. **(C)** Schematic of cells growing inside a microchannel and reaching the channel edge over periods of growth and divisions. **(D)** Mid-slice confocal image (left) and cell wall boundaries before and after ablation (right) of an *rga4Δ* cell, expressing GFP-Psy1, grown inside a microchannel (cyan dotted line) smaller than the standard cell radius in this strain (R_in_). The part of the cell exiting the edge of the channel has a larger radius (R_out_). **(E)** Turgid radius plotted as a function of relaxed radius of *rga4Δ* cells (n=9) grown as in **C-D**, measured both inside (pink) and outside (red) the channel. The line is a linear fit on single cell measurements. **(F)** Elastic strain in the cell wall plotted as a function of cell radius in the same cells as in **E**. Small dots are single cells measurements, and larger dots are means. **(G)** Schematic of the assay to test strain-stiffening hypothesis: cells from each strain are grown in YE5S (YE) in exponential phase, then moved and adapted for 1 h in YE5S+1M sorbitol. Cells from the same sample are then osmotically treated with either YE5S+2M sorbitol (hyperosmotic shock) or YE5S (hypoosmotic shock). **(H)** Relative expansion radius measured as (R_1M_-R_2M_)/R_1M_ for hyper- (dark green), or as (R_YE_-R_1M_)/R_1M_ for hypo- (light green) osmotic treatment in the indicated strains (n=10 cells in each conditions). Error bars are standard deviations. Scale bars 2μm.

As another independent mean to dynamically alter diameter, we used microfabricated PDMS channels. We grew large *rga4Δ* mutant cells into narrow microchannels forcing them to adapt a smaller wild-type like diameter (28, 29). Remarkably, at the channels exit, cells popped out and recovered their normal large diameters in time-scales of few minutes without growing new CW (Fig. S4G-S4I). In a subset of experiments, we could even detect cells with one fatter part out and one thinner part inside the channel (Fig. 4C-4D). This suggests that the stress exerted by turgor pressure on the cell wall is partly compensated by the walls of the microchambers inside the channel, and fully deforms the cell into its normal shape once outside (Fig. 4C-4D and Fig. S4G-S4I). Although lateral wall thickness analysis was not compatible with microchannels, due to unspecific lectin binding to PDMS walls, we could ablate those cells to measure relaxed radii and elastic strains. This analysis yielded again a near conservation of elastic strains, and suggested that expanding walls may rapidly stiffen to limit further expansion (Fig. 4E-4F).

Those last results were indicative of putative strain-stiffening properties of the lateral walls. A signature of materials with such properties is that they compress more than they stretch under an applied mechanical stress of the same magnitude (compressive and tensional, respectively). To assay this, we compared the magnitude of wall deformation in response to hyperosmotic and hypoosmotioc shocks of same amplitudes (30). WT cells were rinsed in 1M sorbitol, and let to adapt and recover their turgid state for 1h. Using flow chambers, we then rapidly rinsed those cells with either 2M sorbitol to create a hyperosmotic shock or normal media for hypoosmotic shocks, and tracked shape changes within less than 2–3 minutes (Fig. 4G). Strikingly, in response to the same pressure difference, cells shrunk significantly more than they inflated (Fig. 4H and Fig. S4J). This differential response was not altered in a *msy2Δ* mutant defective in turgor adaptation to hypoosmotic stress (31), suggesting it primarily arose from strain-stiffening in the wall rather than turgor adaptation. Importantly, this effect was still pronounced in key mutants of the Cell Wall Integrity pathway (32) (Fig. 4H and Fig. S4J), indicating it was mostly independent of downstream wall regulatory signaling, but rather involve passive distortion in the wall polymer network, which stiffen strained walls in a dose-dependent manner.

## Discussion

By combining wall thickness measurements and relaxation assays, we here provide the very first systematic quantitative assessment of CW mechanical properties in the regulation of single cell morphogenesis. We find that the CW is globally or locally stiffer in larger cells or cell portions. This scaling between shape and surface mechanics remains valid among mutants defective in diverse processes such as polarity, signaling and metabolism, and in several species of the *Schizosaccharomyces* clade, suggesting it could represent a broad property of the surface mechanics of walled cells. This effect has dramatic implications on cell size regulation. If the largest fission yeast mutants had similar stiffness as WT cells, they would reach pressurized diameters up to 8–9 µm instead of 6 µm, and *S. japonicus* cells would be typically 2–3X fatter, reaching up diameters of ~15–20µm. Interestingly, we find that wall stiffening in collections of *S. pombe* mutants is mostly associated to an increase in bulk elasticity, suggesting that wall polymer composition or arrangement may be directly or indirectly altered by genetic disruptions. In contrast, other fission yeasts appear to adapt stiffness by thickening their lateral wall at a relatively fixed bulk elasticity. The human pathogen *Cryptococcus neoformans* features titan cells which are typically 10x larger than normal cells and exhibit a specific thick and stiff polymeric capsule (33). The wall of plant-pathogenic fungi can become melanized to support large size increase associated with appressoria formation (33, 34). Modulation of wall surface stiffness may thus be a conserved strategy for walled cells to cope with size variations.

During dynamic shape changes, we provide significant evidence for the existence of gradual wall stiffening which renders stretched CW portions stiffer than relaxed ones. Active lateral wall remodeling, such as in Latrunculin A experiments, may promote stiffening of enlarging cell walls at relatively long time-scales, similar to those of growth. However, physical manipulations in microchambers and with osmotic treatments, support rapid and rather passive strain-stiffening properties of the lateral cell wall. Similar effects have been reported in bacteria and plant cell walls, and proposed to implicate the passive rearrangement of internal wall polymers in the direction of deformation, that render the material stiffer (30, 35). Which particular polymer or cross-link may promote such behavior is an exciting research avenue for the future. We foresee that this mechanical maturation, whether passive or active, could have relevance to normal cell growth, by promoting the stiffening of initially soft CW portions assembled at cell tips, as they move along the cylinder during growth. Thus size-dependent stiffening may confer a unique mechanical benefit to tip growing cells, by limiting lateral size variations, while maintaining a relatively soft CW at growing tips needed for growth (36).

Finally, size-dependent wall stiffening allows for maintaining elastic strains to relatively constant and low values. Measured elastic strains in our assays, do not exceed 25–30%, which is below the estimated failure strain of the CW (at which the CW layer would break open) of ~45% measured in *S. Cerevisiae* (14). Although the composition of the CW in *S. pombe* is different, our findings of a conserved strain well below this failure strain, suggest that the CW will remain mostly intact even when cells are grown to a larger size. Thus, our data support plausible generic mechanical principles of the CW supporting both cell shape and integrity.

## Materials and methods

### Yeast strains, media and genetics

Standard methods for *Schizosaccharomyces pombe* media and genetic manipulations were used (http://www-bcf.usc.edu/~forsburg/). Strains used in this study are listed in supplementary Table 1. Strains indicated with a ‘*’ were derived from the commercially available ‘*S. pombe* Haploid Deletion Mutant Set version 2.0’ strains collection (Bioneer Corporation; http://pombe.bioneer.com). Cells were grown at 25°C in yeast extract plus 5 supplements (YE5S) media.

### Selection of diameter mutants from the KO library

A selection of 62 misshapen mutants was first made by intersecting results from the two reports of visual screens of the KO library (22, 23). Mutants were then transformed with the plasmid pTN509 obtained from the Yeast National Bioresearch Project (Japan), to generate mutant strains expressing GFP-psy1. Alternatively, mutants were crossed with wt strains carrying GFP-psy1, and selected following standard procedures. Strains were visually inspected to confirm the phenotype. To facilitate diameter investigations, only mutants with cylindrical or quasi-cylindrical shape were selected, excluding more complex phenotypes, through a pure qualitative screen. When present, skittle shape phenotype was not totally penetrant, allowing imaging of a subset of cylindrical cells in the population. A final quantification was done, selecting mutants with a larger averaged diameter, or a greater variation (i.e. standard deviation), than the wt. From this screen 18 strains were selected.

### Sample preparation for imaging

In all experiments, cells were pre-labeled in growth media containing 5 μg/ml of labelled lectin from *Griffonia simplicifolia* (alias *Bandeiraea simplicifolia*) *Gs*-IB_4_-Alexafluor647 (Thermo Fisher Scientific) (37). For single-time imaging and laser ablation, cells were usually placed between a glass slide and a coverslip and imaged within 20 min. Time-lapse imaging and osmotic treatments were performed in homemade glass channels built from one 24×50 mm (VWR, Radnor, Pennsylvania, USA) and a 22×22 mm (Thermo Fisher Scientific) coverslip spaced by ~ 250 μm, using double-sided adhesive tape. To make cells adhere to the glass surface, the flow channel was pre-coated with 1 mg/ml poly-L-lysine (Sigma-Aldrich) and 0.1 mg/ml *Gs*-IB_4_ (Sigma-Aldrich), and rinsed with water and YE5S, before pre-labeled cells were placed. Cell diameter confinement was performed by growing cells in microfabricated channels, as described below. Imaging was performed at room temperature (22–26 °C), with controlled humidity (>30%).

### Drug treatments

LatrunculinA (Sigma) was used at a final concentration of 10 μM from a 1000X stock in DMSO. The same amount of DMSO was used for the control. Cells were incubated for 1.5h (or as indicated) at room temperature before imaging for population analysis. For time lapse LatrunculinA (Sigma) treatment, cells were placed in home-made glass channels filled in YE5S+5 mg/ml *Gs*-IB_4_ - Alexafluor647 and a final concentration of 10 μM LatrunculinA from a 1000X stock in DMSO.

### Estimation of relative values of turgor pressure

To estimate pressure values, cells gown in YE5S and pre-labeled with 5 mg/ml *Gs*-IB_4_ - Alexafluor647, were placed in homemade glass channels, imaged, then rinsed with YE5S+5 mg/ml *Gs*-IB_4_ - Alexafluor647 supplemented with different doses of sorbitol and imaged again less than 5 minutes after treatment.

### Dynamic hyper and hypo-osmotic shocks

Cells grown in YE5S were harvested, washed once in YE5S+1M sorbitol, and incubated in the same media for more than 45 min for adaptation. 10 mg/ml *Gs*-IB_4_ - Alexafluor647 was added to the cells during 10 min, and cells were subsequently placed into separate homemade glass channels filled with YE5S+10 mg/ml *Gs*-IB_4_ - Alexafluor647, imaged and rinsed with YE5S for hypoosmotic treatment or with YE5S+2M sorbitol for hyperosmotic treatment. Cells were imaged again less than 3 minutes after treatment.

### Microfabricated chambers

The general design, fabrication and assembly of microchannels to confine fission yeast cells is described in (24). Homothallic *rga4Δ* spores were inserted into the PDMS microchannels, let to germinate for >20h at 25°C in YE5S + Gs-IB4 - Alexafluor647. To allow a better lectin staining the channel was rinsed 2–3 times every 2h before imaging.

### Microscopy

Live-cell imaging was performed on two different inverted spinning-disk confocal microscopes equipped with motorized stages, automatic focus and controlled with MetaMorph^®^ (Microscopy Automation & Image Analysis Software). The first one (Nikon Ti-Eclipse), is equipped with a Yokogawa CSU-X1FW spinning head, and an EM-CCD camera (Hamamatsu), a 100× oil-immersion objective (Nikon CFI Plan Apo DM 100×/1.4 NA) and a 2.5× magnifying lens, yielding a pixel size of 43 nm. The second one (Leica DMI8), is equipped with a Yokogawa CSU-W1 spinning head, and a sCMOS Orca-Flash 4 V2+ (Hamamatsu) a 100× oil-immersion objective (Leica Plan Apo DM 100×oil/1.4 NA), yielding a pixel size of 70nm.

#### Laser ablation assay

The laser ablation set up uses a pulsed 355 nm ultraviolet laser interfaced with an iLas system (Roper Scientific) in the ‘‘Mosquito’’ mode. This allows irradiating at multiple positions in the field with laser spots. This system was mounted on the Nikon confocal spinning disk described above, using a 60x oil-immersion objective (CFI Apochromat 60x Oil λS, 1.4 NA, Nikon) in combination with a 2.5x magnifying lens. The irradiation was performed with a low laser power, in order to minimize bleaching of the lectin labeling, used for shape parameters determination. The irradiation was repeated 3–5 times, until the cell visibly deflated with a clear ejection of cytosolic material observable in bright field, to ensure complete wall relaxation. Cells were imaged before and immediately after ablation, switching to a 100× oil-immersion objective (Nikon CFI Plan Apo DM 100×/1.4 NA) and a 2.5× magnifying lens, yielding a pixel size of 43 nm. The full process, including the first image acquisition, laser ablation and the second image after ablation, was typically performed in less than 2 minutes.

Imaging for chromatic registration was performed as in (16), by imaging a slide containing a solution of 0.2 µm TetraSpeck™ microspheres (Thermofisher), and generating images of the same bead at different positions in the field of view to create an ordered array.

Details for image analysis scripts and general methods are reported in supplemental informations.

### Extraction of bulak elastic moduli from wall relaxation assays

#### Subcellular estimation of Y/P

Values of bulk modulus divided by turgor pressure were estimated by the force balance equation, as described in (11), using the variations in cell shape before and after laser ablation. To this, we implemented local measurements of wall thickness, measured in the inflated state (before ablation), obtaining local values of wall thickness in each cell analyzed. For cell side the force balances on the CW yields:

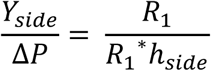
where 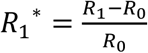, with R_0_ the deflated radius on cell sides. At cell tips, force balance reads:

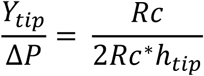
where 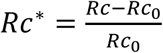, with Rc_0_ the deflated curvature radius at cell tips.

Old tips and new tips were analyzed separately.

#### Subcellular estimation of Y/P with a simulation method

When local cell shapes were not perfectly cylindrical, for instance to analyze bulk moduli of the birth scars (Fig 1D), we derived values for the bulk modulus using a computational approach. The same approach was used to confirm experimental estimations in wt, and skittle-like cells. This computational method yielded near-exact values as the aforementioned analytical formula for rod and rod-like parts of the cells (Fig S1C and Fig. S3D). This method was similar than the one presented in (10, 16), and details are reported in supplemental methods.

#### Estimation of relative values of turgor pressure

Turgor pressure was calculated as previously described in (11). The external osmolarity at which the cell wall reaches its relaxed state was estimated from a comparison with laser ablation data, considering this last as the fully relaxed wall state (Fig. S2B). The same values for sorbitol, glycerol and YE osmolarity, molarity to molality conversion coefficients, and cell inaccessible volume fractions were used as in (11). Volumes before (V_1_) and after (V_0_) ablation were measured at the population of each mutant, considering the central part of the cell as a cylinder and the two ends as hemispheres.

## Acknowledgments

The authors acknowledge Y. Sanchez, T. Toda, P. Perez and N. Rhind for sharing material. We thank S. Dmitrieff, L. Chevallier, A. Haupt and other members of the Minc laboratory. We acknowledge the ImagoSeine facility, a member of France BioImaging (ANR-10-INSB-04). This project was supported by the CNRS and grants from the FP7 CIG program, ITN “FungiBrain”, the ANR (“CellSize”) and the European Research Council (CoG Forcaster no. 647073).

