## Supplemental Material for "Systematic Mapping of cell Wall Mechanics in the Regulation of Cell Morphogenesis"

### SUPPORTING INFORMATION

This files contains supplemental methods, supplemental references, 2 supplemental tables, 4 supplemental figures and figure legends

### Supplemental Methods

#### Image analysis:

*Cell Wall thickness computation:* Wall thickness measurement was performed as in (1), including chromatic shift registration and correction. Images of cells labeled for the plasma membrane (GFP-Psy1) and cell wall surface (*Gs*-$IB_4$-Alexafluor647) were analyzed. For measurement in other fission yeasts, the membrane labelling was provided using FM-464. For this, cells were incubated for 10min in the dye at a final concentration of 5µg/ml from a 100X stock in DMSO. This provided sharp membrane labelling on cell sides, but did not allow to map thickness at cell tips. From those signals, a first binarization defined the most external cell boundary, on which perpendicular lines were defined at each pixel. Because some of the mutants showed a variable intracellular background, plausibly due to partial defects in Psy1 delivery to plasma membrane or increased auto-fluorescence, we systematically implemented a correction script to extract the true signal peak by using an analytical expression of the convoluted intensity profile (1, 2).

*Cell Shape and local Wall thickness measurements:* For each cell, wall boundaries were obtained by binarization of mid-slice confocal fluorescent images of lectin-Alexafluor647 label, before and after laser ablation. For some cells, the results were manually updated, to correct for local signal bleaching after laser ablation. The long axis of the cell was automatically detected and defined as cell length. Old and new ends were defined by the user, and local radii of curvature ($R_{c1-0}$) were measured by fitting local tips with a circle using a portion of cell surface of 3.44 µm around the center of the tip; the same section was used to measure tip wall thickness. The two scar bulges localization was defined by the user, scar radius was automatically measured as the distance between them, and thickness computed as a mean of values on a distance of 0.430 µm around the selected center. Cell sides limits (single or multiple) were defined by the user, and the radius of the cell ($R_{1-0}$) was measured as the mean of all diameters on the selected region, as well as side thickness. Birth scars were sometimes superimposed on the selected side region, as in multi-scarred cells. In such case, the scar region (0.430 µm around the selected center) was automatically excluded to compute side parameters.

The analysis of shape changes during osmotic treatments, were computed from the lectin signal using ImageJ. The diameter was calculated as an average of 5 measurements along the cylindrical part of the cell.

*Subcellular estimation of Y/P with a simulation method:*

The estimation procedure of cell mechanical parameters (the scars in Figure 1D, the old end, new end and cell sides in Figure S1C) is achieved using the following procedure: The plasmolized contour of the cell is swollen with an internal pressure of 1MPa supposing a one-parameter family of Young-modulus distribution, the parameter fitting the best the turgid contour is retained to calculate the estimate. The chosen one-parameter families of Young modulus distributions are the following one:

- For the estimation of the scar modulus Figure 1D, the distribution is given at the curvilinear abscissa x (The 0 of the curvilinear abscissa being taken at the oe and the abscissa being expressed in μm) by the formula:

$$Y(x)=(Y_{oe}+((Y_{side}-Y_{oe})*(\text{sigmoid}(x,(x_{oe}+w_{tip}+x_{side\_center})/2,5/(x_{side\_centre}/2-x_{oe}-w_{tip}))))+((Y_{scar}-Y_{side})*(\text{sigmoid}(x,(x_{scar}-w_{scar}+x_{side\_centre})/2,10/(x_{side\_centre}-x_{scar}-w_{scar}))))+((Y_{ne}-Y_{scar})*(\text{sigmoid}(x,(x_{scar}+w_{scar}+x_{ne}-w_{tip})/2,5/(x_{ne}-w_{tip}-x_{scar}-w_{scar}))))).$$

$Y_{oe}$,$Y_{side}$,$Y_{ne}$ are estimated independently using analytical formula for spherical, and cylindrical pressure vessel; $Y_{scar}$ is the sole free-parameter.

- In Figure S1C, the side modulus is estimated by running simulations with: Y(x)= Yside. The $Y_{sidesim}/P$ estimated through simulations is 1.07±0.025(N=27) times smaller than $Y_{side}/P$ estimated using the analytical formula for a cylindrical pressure vessel.

- In Figure S1C, the oe and ne modulus are estimated by running simulations with the following Young modulus distribution:

$$Y(x)=(Y_{oesim}+((Y_{side}-Y_{oesim})*(\text{sigmoid}(x,(x_{oe}+w_{tip}+x_{side\_center})/2,5/(x_{side\_centre}/2-x_{oe}-w_{tip}))))+((Y_{nesim}-Y_{side})*(\text{sigmoid}(x,(x_{ne}\,w_{tip}+x_{side\_centre})/2,10/(x_{ne}-w_{tip}-x_{side\_center}))))).$$

$Y_{oesim}=(1-t)*Y_{oe}+t*Y_{side}$, $Y_{nesim}=(1-t)*Y_{ne}+t*Y_{side}$. $Y_{oe}$,$Y_{side}$,$Y_{ne}$ are estimated independently using analytical formula for spherical, and cylindrical pressure vessel. For each cell, the best fitting parameter t is retained to estimate $Y_{oesim}$ and $Y_{nesim}$. The t estimation is: 0.3704±0.13(N=27).

In each of the distribution, $w_{tip}$= 1.7 µm corresponds to the tip half-width and $w_{scar}$= 0.21 µm corresponds to the scar half-width.

The distribution of Young modulus in the central zone of the cell in Figure S3E is calculated using the thickness profile and the diameter of the cell through the formula of the linear regressions from Figure3B for *myo1Δ* and *ubr1Δ* supposing P=1MPa. The distributions are connected at both tips to the analytical estimations of the moduli by fitting a second order polynomial function between the cell wall portion closest from the tip and the tip itself. For *myo1Δ*, the simulated swelling in FigureS3E corresponds to P=1MPa and for *ubr1Δ*, the simulated swelling FigureS3E corresponds to P=1.3MPa (the Young modulus estimated Figure3B is slightly too stiff).

*GFP-Bgs4 concentration measurements:* To analyze the localization of polar factors we used the same approach as in (1). Cells expressing GFP-Bgs4 were labeled with *Gs*-$IB_4$ -Alexafluor647. The cell was first segmented by using the signal from the lectin-labelled cell wall, in order to extract the whole-cell contour. Fluorescent signals of interest were then extracted from fluorescent images by using a mask based on corresponding sub-regions. Signals were corrected for the background signal:

$$I_{tip} = I_{tip\ raw} - I_{bg}$$

$$I_{cell\ contour} = I_{cell\ contour\ \ raw} - I_{bg}$$

Normalized tip signal is the ratio between the averaged intensity at the two tips divided by the intensity along the full membrane.

$$Norm - Bgs4\ Tip\ Signal = \frac{I_{tip}}{I_{cell\ contour}}$$

*Cell 3D reconstruction:* To analyze cell shapes in microchannel experiments, z-stacks of the membrane labeling GFP-Psy1 covering the whole depth of the cell were acquired with a z-interval of 200 nm. Images were segmented using the Limeseg plugin for Fiji (3), which converts the stack into a 3D surface mesh. This reconstructed cell shape was analyzed using a custom Matlab script available online at https://github.com/SergeDmi/analyze_pombe. A closed curve was fitted to the points of the central slice using a spline-fitting tool adapted from (4); the perimeter was determined as the length of this curve.

| Gene ID | GO biological processes | | | | | |
|---|---|---|---|---|---|---|
| ***scd2*** | signaling | establishment or maintenance of cell polarity | conjugation with cellular fusion | | | |
| ***myo52*** | mitotic cytokinesis | actin cytoskeleton organization | establishment or maintenance of cell polarity | | | |
| ***coq10*** | cofactor metabolic process | | | | | |
| ***kin1*** | mitotic cytokinesis | membrane organization | actin cytoskeleton organization | establishment or maintenance of cell polarity | | |
| ***pub1*** | protein catabolic process | transmembrane transport | protein modification by small protein conjugation or removal | | | |
| ***mae2*** | generation of precursor metabolites and energy | | | | | |
| ***vps1*** | peroxisome organization | membrane organization | vesicle-mediated transport | | | |
| ***dad1*** | meiotic nuclear division | mitotic sister chromatid segregation | | | | |
| ***yaf9*** | DNA repair | regulation of transcription, DNA-templated | chromatin organization | transcription, DNA-templated | | |
| ***plc1*** | signaling | lipid metabolic process | | | | |
| ***gpa1*** | ascospore formation | signaling | conjugation with cellular fusion | | | |
| ***myo1*** | membrane organization | protein-containing complex assembly | actin cytoskeleton organization | vesicle-mediated transport | establishment or maintenance of cell polarity | |
| ***efc25*** | signaling | establishment or maintenance of cell polarity | | | | |
| ***ubr1*** | signaling | protein catabolic process | regulation of transcription, DNA-templated | protein modification by small protein conjugation or removal | transcription, DNA-templated | |
| ***rdp1*** | regulation of transcription, DNA-templated | meiotic nuclear division | chromatin organization | transcription, DNA-templated | | |
| ***rga2*** | signaling | | | | | |
| ***rga4*** | signaling | actin cytoskeleton organization | establishment or maintenance of cell polarity | | | |
| ***mtq2*** | cytoplasmic translation | | | | | |
| ***wsc1*** | signaling | | | | | |
| ***mtl2*** | signaling | | | | | |
| ***rgf2*** | cell wall organization or biogenesis | signaling | | | | |
| ***rgf1*** | signaling | carbohydrate metabolic process | cell wall organization or biogenesis | | | |
| ***pmk1*** | metal ion homeostasis | signaling | transmembrane transport | regulation of transcription, DNA-templated | transcription, DNA-templated | cell wall organization or biogenesis |
| ***pck2*** | signaling | carbohydrate metabolic process | establishment or maintenance of cell polarity | cell wall organization or biogenesis | | |
| ***agn1*** | carbohydrate metabolic process | conjugation with cellular fusion | cell wall organization or biogenesis | | | |
| ***omg1*** | carbohydrate metabolic process | conjugation with cellular fusion | cell wall organization or biogenesis | | | |

**Table S1:** Gene functions for our set of mutants assigned from the Gene Ontology classification (https://www.pombase.org/)

| | | |
|---|---|---|
| h+ *GFP-psy1:ade bgs4::ura4 RFP-bgs4-Leu* (*leu1-32 ura4-D18 ade6*) | This study | VD57 |
| h- *GFP-psy1:ade bgs4::ura4 RFP-bgs4-Leu* (*leu1-32 ura4-D18 ade6*) | This study | VD58 |
| h+ *bgs4::ura4 RFP-bgs4-Leu* (*leu1-32 ura4-D18 ade6-M216*) | This study | NM387 |
| h- *rga2::KanMX bgs4::ura4 RFP-bgs4-Leu GFP-psy1:ade* (*leu1-32 ura4-D18*) | This study | VD72 |
| h- *rga4::KanMX bgs4::ura4 RFP-bgs4-Leu GFP-psy1:ade* (*leu1-32 ura4-D18*) | This study | VD81 |
| *h+ wsc1::KanMX GFP-psy1:ade bgs4::ura4 RFP-bgs4:leu* (*leu1-32 ura4-D18 ade6*) | This study | VD158 |
| h- *mtl2::KanMX GFP-psy1:ade bgs4::ura4 RFP-bgs4:leu* (*leu1-32 ura4-D18 ade6*) | This study | VD160 |
| h- *pck2::leu GFP-psy1:ade* | This study | VD109 |
| h+ *rgf2::ura GFP-psy1:ade* (*leu1-32 ura4-D18 ade6*) | This study | VD164 |
| h+ *rgf1::KanMX GFP-psy1:ade* (*leu1-32 ura4-D18 ade6*) | This study | VD167 |
| h+ *pmk1:ura GFP-psy1:leu* (*leu1-32 ura4-D18*) | This study | VD173 |
| h+ *scd2::KanMX CRIB-tdTomato:ura Leu:GFP-psy1 (ura4-D18 leu1-32)* | This study | VD97 |
| h+ *myo52::ura CRIB-tdTomato:ura Leu:GFP-psy1 (ura4-D18 leu1-32)* | This study | VD127 |
| h+ *coq10::KanMX GFP-psy1:ade (ade6-M216 ura4-D18 leu1-32)* | This study | HG77* |
| h+ *kin1::KanMX GFP-psy1:ade (ade6-M216 ura4-D18 leu1-32)* | This study | HG101* |
| h+ *pub1::KanMX GFP-psy1:ade (ade6-M216 ura4-D18 leu1-32)* | This study | HG72* |
| h+ *mae2::KanMX GFP-psy1:ade (ade6-M216 ura4-D18 leu1-32)* | This study | HG74* |
| h+ *vps1::KanMX GFP-psy1:ade (ade6-M216 ura4-D18 leu1-32)* | This study | HG31* |
| h+ *dad1::KanMX GFP-psy1:ade (ade6-M216 ura4-D18 leu1-32)* | This study | HG76* |
| h+ *yaf9::KanMX GFP-psy1:ade (ade6-M216 ura4-D18 leu1-32)* | This study | HG103* |
| h+ *plc1::KanMX GFP-psy1:ade (ade6-M216 ura4-D18 leu1-32)* | This study | HG104* |
| h+ *gpa1::KanMX GFP-psy1:ade (ade6-M216 ura4-D18 leu1-32)* | This study | HG106* |
| h+ *myo1::KanMX: GFP-psy1:ade (ade6-M216 ura4-D18 leu1-32)* | This study | HG27* |
| h+ *efc25::KanMX GFP-psy1:ade (ade6-M216 ura4-D18 leu1-32)* | This study | HG29* |
| h+ *ubr1::KanMX GFP-psy1:ade (ade6-M216 ura4-D18 leu1-32)* | This study | HG33* |
| h+ *rdp1::KanMX GFP-psy1:ade (ade6-M216 ura4-D18 leu1-32)* | This study | HG35* |
| h+ *mtq2::KanMX GFP-psy1:ade (ade6-M216 ura4-D18 leu1-32)* | This study | HG116* |
| h+ *agn1:KanMX GFP-psy1:ade (ade6-M216 ura4-D18 leu1-32)* | This study | HG75* |
| h+ *omg1:KanMX GFP-psy1:ade (ade6-M216 ura4-D18 leu1-32)* | This study | HG121* |
| h90 rga4::KanMX ade6<<GFP-psy1 bgs4::ura4 RFP-bgs4-Leu | This study | VD162 |
| h+ msy2::KanMX *(ade6-M216 ura4-D18 leu1-32)* | This study | AH 233* |
| *Schizosaccharomyces japonicus, wild-type, homothallic* | YGRC | VD210 |
| *Schizosaccharomyces octosporus*, wild-type, homothallic | YGRC | VD211 |
| *Schizosaccharomyces cryophilus*, wild-type, homothallic | Nick Rhind | VD212 |

**Table S2.** Yeast strains used in this study

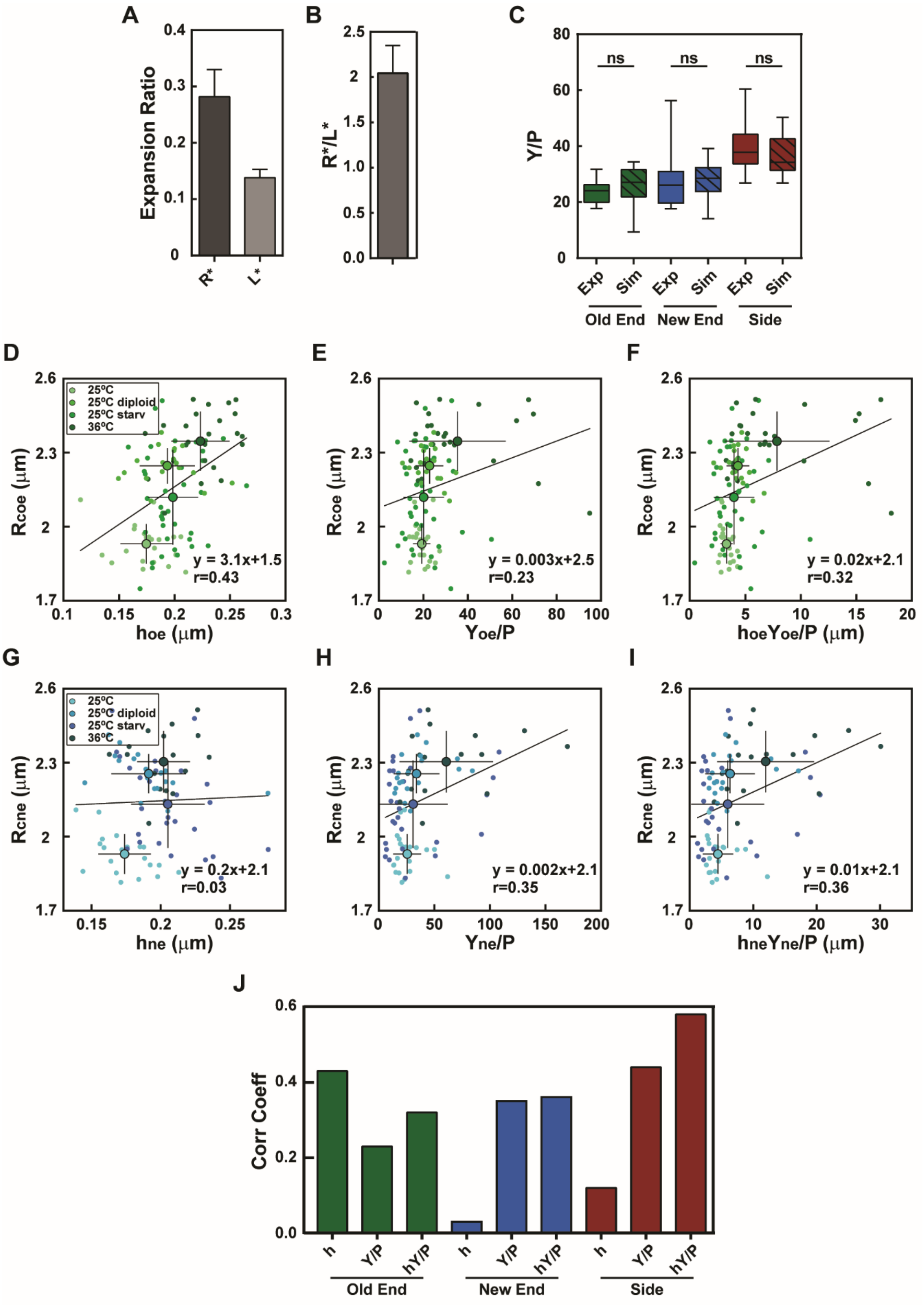

A
Expansion Ratio
R*
L*
B
R*/L*
C
Y/P
ns
Exp
Sim
Old End
New End
Side
D
E
F
Rcoe (µm)
hoe (µm)
Yoe/P
hoeYoe/P (µm)
25ºC
25ºC diploid
25ºC starv
36ºC
y = 3.1x+1.5
r=0.43
y = 0.003x+2.5
r=0.23
y = 0.02x+2.1
r=0.32
G
H
I
Rcne (µm)
hne (µm)
Yne/P
hneYne/P (µm)
y = 0.2x+2.1
r=0.03
y = 0.002x+2.1
r=0.35
y = 0.01x+2.1
r=0.36
J
Corr Coeff
h
Y/P
hY/P

**Figure S1. Local cell wall mechanics and cell shape in fission yeast.**

**(A)** Relative expansion radius (measured as $(R_1-R_0)/R_0$) and relative expansion length (measured as $(L_1-L_0)/L_0$) of wt cells after laser ablation (n=99). **(B)** Ratio between the two expansion ratios calculated for each cell from **A** (n=99). **(C)** Y/P values computed experimentally (plain) or simulated (striped) at old end, new end and sides, on the same population of WT cells (n=27). Whisker plots represent median and full data set range. **(D-F)** Radius of curvature at the old end ($R_{oe}$), plotted as a function of old end wall thickness ($h_{oe}$), old end bulk modulus divided by P ($Y_{oe}/P$) or surface modulus divided by P ($Y_{oe}h_{oe}/P$), in the same conditions as in **Fig. 1G** (for 25$^0$C n=21, 25$^0$C diploid n= 18, 25$^0$C starv n=39, 36$^0$C n=25). **(G-I)** Radius of curvature at the new end ($R_{ne}$), plotted as a function of new end wall thickness ($h_{ne}$), new end bulk modulus divided by P ($Y_{ne}/P$) or surface modulus divided by P ($Y_{ne}h_{ne}/P$), in the same conditions as in **Fig. 1G** (for 25$^0$C n=21, 25$^0$C diploid n= 19, 25$^0$C starv n=29, 36$^0$C n=17). For **D-I**, small dots correspond to single cells measurements, and the larger dot is the average. The line is a linear fit of single cell measurements. Error bars are standard deviations. **(J)** Pearson correlation coefficients, between the local radius and wall thickness (h), bulk modulus divided by pressure Y/P or surface modulus divided by pressure (hY/P), in different parts of the cell as indicated. Those values correspond to those presented in **D-E** and in **Fig. 1 H-J.**

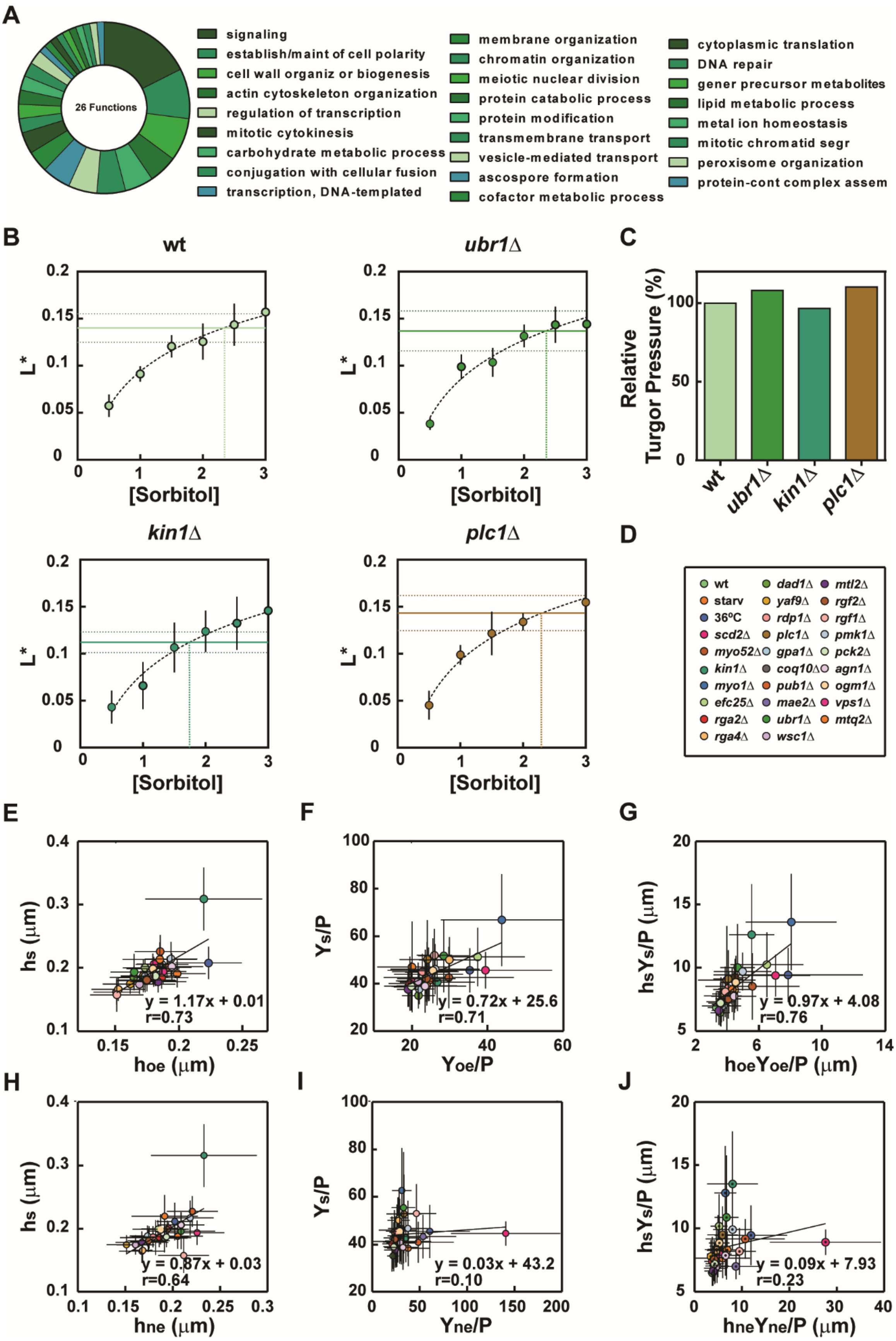
A
26 Functions
signaling
establish/maint of cell polarity
cell wall organiz or biogenesis
actin cytoskeleton organization
regulation of transcription
mitotic cytokinesis
carbohydrate metabolic process
conjugation with cellular fusion
transcription, DNA-templated
membrane organization
chromatin organization
meiotic nuclear division
protein catabolic process
protein modification
transmembrane transport
vesicle-mediated transport
ascospore formation
cofactor metabolic process
cytoplasmic translation
DNA repair
gener precursor metabolites
lipid metabolic process
metal ion homeostasis
mitotic chromatid segr
peroxisome organization
protein-cont complex assem
B
wt
ubr1Δ
kin1Δ
plc1Δ
L*
[Sorbitol]
C
Relative Turgor Pressure (%)
D
wt
starv
36ºC
scd2Δ
myo52Δ
kin1Δ
myo1Δ
efc25Δ
rga2Δ
rga4Δ
dad1Δ
yaf9Δ
rdp1Δ
plc1Δ
gpa1Δ
coq10Δ
pub1Δ
mae2Δ
ubr1Δ
wsc1Δ
mtl2Δ
rgf2Δ
rgf1Δ
pmk1Δ
pck2Δ
agn1Δ
ogm1Δ
vps1Δ
mtq2Δ
E
hs (μm)
hoe (μm)
y = 1.17x + 0.01
r=0.73
F
Ys/P
Yoe/P
y = 0.72x + 25.6
r=0.71
G
hsYs/P (μm)
hoeYoe/P (μm)
y = 0.97x + 4.08
r=0.76
H
hs (μm)
hne (μm)
y = 0.87x + 0.03
r=0.64
I
Ys/P
Yne/P
y = 0.03x + 43.2
r=0.10
J
hsYs/P (μm)
hneYne/P (μm)
y = 0.09x + 7.93
r=0.23

**Figure S2. Classification of diameter defective mutants, variations in turgor values between mutants, and correlations between tips and lateral cell wall mechanics.**

**(A)** Pie chart of Gene Ontology classification of the mutants used in the screen presented in **Fig. 2**. The list starts with the most abundant and follows in a clockwise direction. **(B)** Relative expansion length of cells from the indicated strains, grown in YE5S and treated with different sorbitol concentrations in YE5S. Dots represent an average of a treated population (n>10). L* is measured as $(L_1-L_{osm})/L_{osm}$ ) where $L_{osm}$ is the length acquired after osmotic treatment. The horizontal plain line represents an average of the relative expansion length obtained in the same strain using laser ablation ($L_{osm}=L_0$), horizontal dotted lines indicate the corresponding standard deviation. Vertical line starts from the intersection between averaged L* and logarithmic fit, and ends in the intersection with the *x* axis **(C)** Turgor pressure of indicated strains, relative to wt turgor pressure, calculated as $(P_{strain}/P_{wt})$*100. **(D)** Legend for the conditions or mutants used in **E-J.** **(E-G)** CW thickness, bulk elastic modulus divided by pressure, and surface modulus divided by pressure on cell sides plotted as a function of those at old ends. **(H-J)** CW thickness, bulk elastic modulus divided by pressure, and surface modulus divided by pressure on cell sides plotted as a function of those at new ends. r values correspond to Pearson correlation coefficients. Error bars are standard deviations.

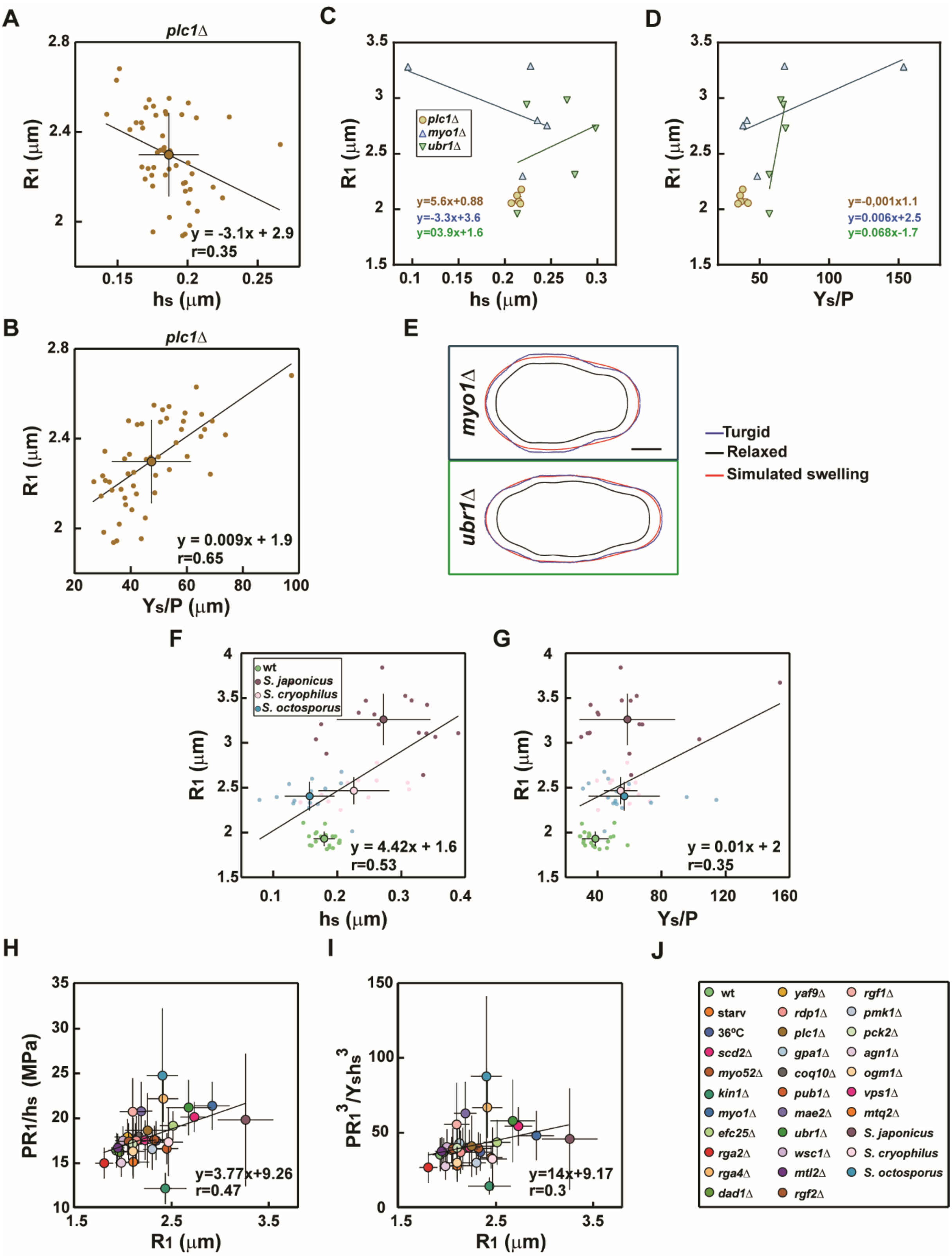
A
plc1Δ
R1 (µm)
hs (µm)
y = -3.1x + 2.9
r=0.35
C
R1 (µm)
hs (µm)
plc1Δ
myo1Δ
ubr1Δ
y=5.6x+0.88
y=-3.3x+3.6
y=03.9x+1.6
D
R1 (µm)
Ys/P
y=-0,001x1.1
y=0.006x+2.5
y=0.068x-1.7
B
plc1Δ
R1 (µm)
Ys/P (µm)
y = 0.009x + 1.9
r=0.65
E
myo1Δ
ubr1Δ
Turgid
Relaxed
Simulated swelling
F
wt
S. japonicus
S. cryophilus
S. octosporus
R1 (µm)
hs (µm)
y = 4.42x + 1.6
r=0.53
G
R1 (µm)
Ys/P
y = 0.01x + 2
r=0.35
H
PR1/hs (MPa)
R1 (µm)
y=3.77x+9.26
r=0.47
I
PR1^3/Yshs^3
R1 (µm)
y=14x+9.17
r=0.3
J
wt
starv
36ºC
scd2Δ
myo52Δ
kin1Δ
myo1Δ
efc25Δ
rga2Δ
rga4Δ
dad1Δ
yaf9Δ
rdp1Δ
plc1Δ
gpa1Δ
coq10Δ
pub1Δ
mae2Δ
ubr1Δ
wsc1Δ
mtl2Δ
rgf2Δ
rgf1Δ
pmk1Δ
pck2Δ
agn1Δ
ogm1Δ
vps1Δ
mtq2Δ
S. japonicus
S. cryophilus
S. octosporus

**Figure S3. Evolution of key mechanical values in the CW as a function of cell radius.** **(A-B)** Cell radius ($R_1$) plotted as a function of side wall thickness ($h_s$, **A**) and bulk modulus divided by P ($Y_s$/P, **B**) within a population of *plc1Δ* (n=50). **(C-D)** Local radius ($R_1$), plotted as a function of the corresponding side wall thickness ($h_s$, **C**) and bulk modulus divided by P ($Y_s$/P, **D**) in 5 different parts along the side of the same cell, for the cells represented in **Fig. 3B**. **(E)** Simulation of cell swelling in skittle cells of indicated strains, obtained by using values of local surface moduli estimated experimentally as in **C-D** and **Fig. 3B**. In black and blue: measured relaxed and turgid cell boundary, in red: simulated cell boundary. **(F)** Cell radius ($R_1$) plotted as a function of side wall thickness (h), **(G)** and side bulk modulus divided by P ($Y_s$/P) in *S. pombe* (n=21), S. *japonicus* (n=17), *S. octosporus* (n=14), and *S. cryophylus* (n=12). **(H)** Elastic stress in the cell wall ($PR_1/h_s$) plotted as a function of cell radius and **(I)** energy corresponding to cell wall bending normalized to the energy of pressure ($PR_1/h_sY_s$) plotted as a function of cell radius **(J)** Legend of conditions or mutants used in **H-I.** Where present, small dots are single cells measurements, and the big dot is the average. In A-D,F-G the line is a linear fit of single cell measurements. In H-I the line is a linear fit of averages. r values corresponds to Pearson correlation coefficients. Error bars represent standard deviations. Scale bar is 2µm.

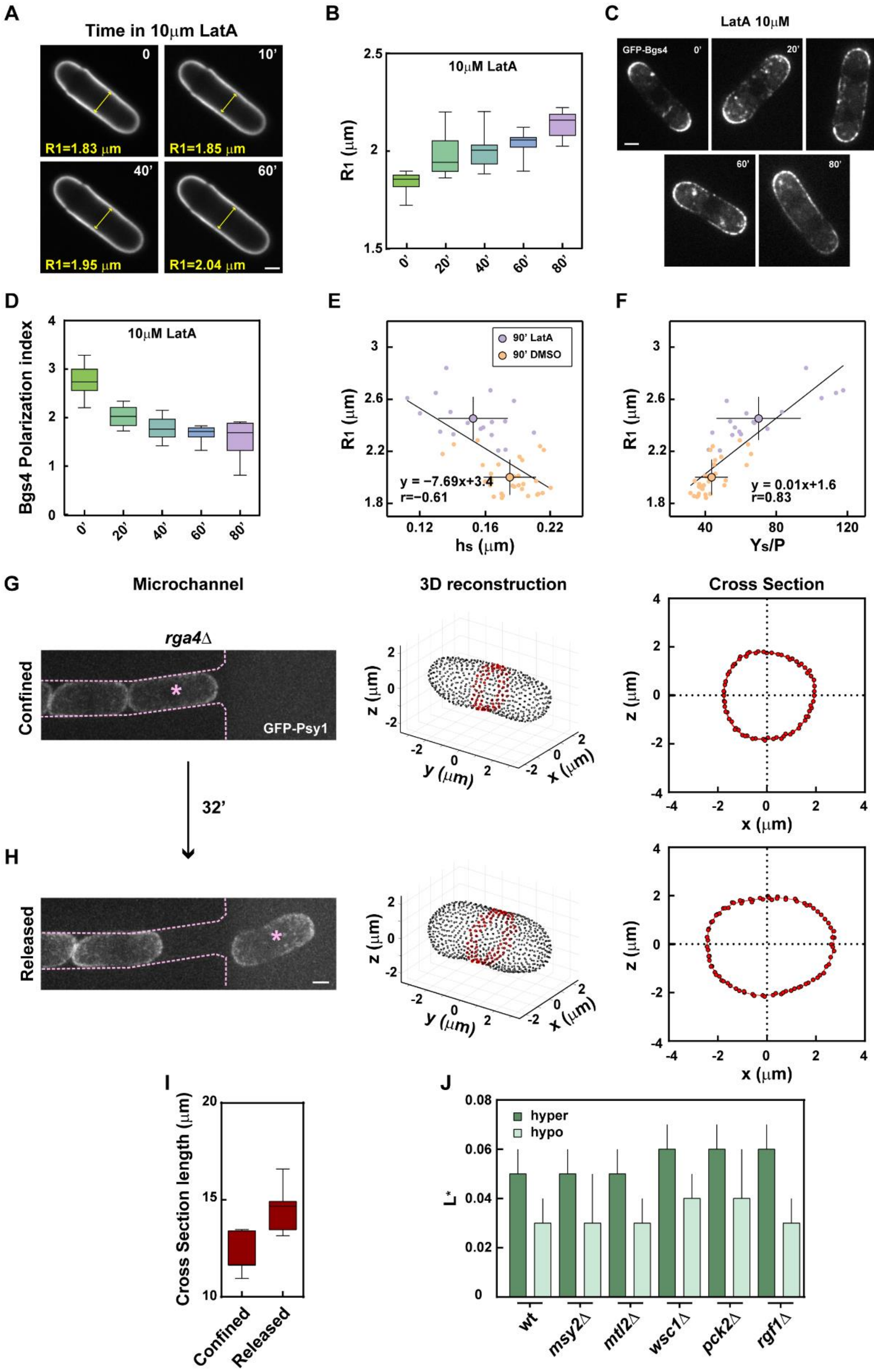
A
Time in 10μm LatA
R1=1.83 μm
R1=1.85 μm
R1=1.95 μm
R1=2.04 μm
B
10μM LatA
R1 (μm)
C
LatA 10μM
GFP-Bgs4
D
Bgs4 Polarization index
10μM LatA
E
90' LatA
90' DMSO
y = −7.69x+3.4
r=−0.61
hs (μm)
F
y = 0.01x+1.6
r=0.83
Ys/P
G
Microchannel
3D reconstruction
Cross Section
rga4Δ
Confined
GFP-Psy1
32'
H
Released
I
Cross Section length (μm)
Confined
Released
J
hyper
hypo
L*
wt
msy2Δ
mtl2Δ
wsc1Δ
pck2Δ
rgf1Δ

**Figure S4. Evolution of cell diameter and CW mechanics upon dynamic shape manipulations.**

**(A)** Time-lapse of shape changes in a wt cell after treatment with 10μM Latrunculin A. The cell is labeled with lectin *Gs*-Alexafluor647. **(B)** Quantification of cell radius ($R_1$) evolution as a function of time. **(C)** Typical GFP-Bgs4 localization after treatment with 10μM Latrunculin A. **(D)** Quantification of GFP-Bgs4 polarity (tip signal divided by full cell contour signal) in cells treated as in **A**. **(E)** Mean cell radius ($R_1$), plotted as a function of side wall thickness ($h_s$), and **(F)** side bulk modulus ($Y_s/P$) in wt cells treated for 1.5 hr with DMSO (n=30) or 10μM LatrunculinA (n=18). Small dots are single cells measurements, and larger dots are averages. Lines are linear fits on single cell measurements. **(G-H)** (Left) Maximum projection of 15 slices of *rga4Δ* cells expressing GFP-psy1 and grown inside a microchannel. The cell indicated by the asterisk is confined in (G) and released in (H). Note the rapid change in cell diameter. 3D reconstruction (Center) and cross section (Right) of the cell marked with an asterisk on the left before and after exiting the microchannel. **(I)** Cell perimeter in a population of *rga4Δ* cells grown in microchannels before and after release. (n=7) **(J)** Relative expansion length measured as $(L_{1M}-L_{2M})/L_{1M}$ for hyper (dark green), or as $(L_{YE}- L_{1M})/L_{YE}$ for hypo (light green) osmotic treatment in cells treated as in **Fig. 4 H**, in the indicated strains (n=10 in each conditions). Error bars are standard deviations, scale bars 2μm.
